# Convergence and constraint in glucosinolate evolution across the Brassicaceae

**DOI:** 10.1101/2025.04.23.650103

**Authors:** Amanda Agosto Ramos, Kevin A. Bird, Annanya Jain, Gabriel Phillip, Odinaka Okegbe, Lucy Holland, Daniel Kliebenstein

**Affiliations:** Department of Plant Sciences, University of California, Davis One Shields Ave, Davis, CA, 95616 USA; Plant Biology Graduate Group, University of California, Davis One Shields Ave, Davis, CA, 95616 USA; Genetic and Genomics Graduate Group, University of California, Davis One Shields Ave, Davis, CA, 95616 USA

## Abstract

Diversity in plant specialized metabolites plays critical roles in plant-environment interactions. In longer evolutionary scales, e.g. between families or orders, this diversity arises from whole-genome and tandem duplication events. Less is known about the evolutionary patterns shaping chemical diversity at shorter scales, e.g. within a family. Utilizing the aliphatic glucosinolate pathway we explored how the terminal structural modification enzyme GSL-OH evolved across the *Brassicaceae* and the genomic processes that control presence-absence variation of its products (R)-2-hydroxy-but-3-enyl and (S)-2-hydroxy-but-3-enyl. We implemented a phylo-functional approach where we functionally validated GSL-OH orthologs across the Brassicaceae and used that information to map the genomic origin and trajectory of the loci. This uncovered a complex mechanism involving at least three ancestral loci with extensive gene loss across all species creating unequal retention across the phylogenetic relationships. Convergent evolution in enantiomeric specificity was observed where several independent species had tandem duplicates diverged towards producing the R or S enantiomers. To explore potential biological differences between the enantiomers, we performed *Trichoplusia ni* insect assays and *Botrytis cinerea* detached leaf assays. We found that plants with the S enantiomer were more susceptible to *B. cinerea* infection while plants with the R enantiomer seemed more susceptible to *T. ni* herbivory. This variation in activity between the precursor and different enantiomeric products may shape the recurrent evolution of their production.

## Introduction

Plant specialized metabolites mediate the myriad of plant-biotic and abiotic interactions by having a broad range of biological functions and mechanisms (Weng, J., et al., 2021). These potential functions have recently expanded to roles as internal signaling compounds (Katz et al., 2015), and storage molecules (Petersen et al., 2002), that can be recycled and re-incorporated to central metabolism (Sugiyama et al., 2021). The tasks plant specialized metabolites fulfill often involve strong and variable evolutionary pressures that contribute to shaping the wide range of specialized metabolite structural diversity. These roles have led to an increased interest in identifying the pathways that generate this chemical diversity pathways or investigating the structure/function relationships in key enzymes across evolution (Weng, J., et al., 2021). Less work has queried how well-known biosynthetic pathways evolutionarily change across closely related species.

As key pathways are identified in model species there is a growing appreciation of the need to move beyond model species to understand how specialized metabolite pathways evolve to create diversity. Chemical analysis has shown extensive shifts in plant specialized metabolic diversity across all phylogenetic levels from order, to family, genus, between and within species level. This work has developed a cornerstone model suggesting that family/genus level diversity is often a result of whole genome duplication events that enable a duplicate gene to undergo neofunctionalization and create new enzymes that form new pathways (Hofberger et al., 2013). These new enzymes can create a core structure that is then chemically modified to create extensive chemical diversity at the family to species level. At even shorter time scales, e.g. comparing closely related species or within species variation, specialized metabolites show extensive presence/absence variation with complex patterns that make it difficult to assess if the underlying processes are parallel or convergent evolution or rampant independent gene loss (Ono and Murata, 2023). This complicates the ability to understand the contribution of whole genome duplications versus tandem/distal duplications in this range of specialized metabolite variation. Identifying the processes facilitating broader diversity of specialized metabolites from the species to family level is needed to develop a deeper model of how diversity in specialized metabolism is generated.

Glucosinolates, a family of specialized metabolites unique to the Brassicales, are a model system to investigate the genetic and genomic processes creating chemical diversity (Hofberger et al., 2013). With 137 unique glucosinolate structures identified thus far, the pathway is considered relatively simpler and younger than other pathways that synthesize 1000s of different structures across numerous orders (Blažević et al., 2020). Glucosinolates are grouped by the amino acid used to create the core structure, with the methionine-derived glucosinolate pathway in the Brassicaceae family having facilitated a rapid expansion of structural diversity (Bird et al., 2025). This aliphatic glucosinolate structural diversity in *A. thaliana* and Brassica spp is caused by three main loci: MAMs, AOPs, and GSL-OH (Kliebenstein and Cacho, 2016). Across the Brassicaceae, these genes show epistatic presence-absence variation where production of the final 2-hydroxy-but-3-enyl (2HB3) requires a functional GSL-OH, AOP2 and 4C capable MAM. Modular variation in these enzymes controls structural diversity by determining chain length ranging from 2 to 11 carbons, and which chemical modifications occur on the side chain. Each of these enzymes/loci contain extensive and complex presence/absence patterns of biosynthesis genes in *A. thaliana* and domesticated Brassica spp that complicate parsimony interpretations. For example, initial function-to-phenotype analysis of chain elongation in *Brassica* and *Arabidopsis* identified three MAM enzymes (1/2/3) their presence or absence appeared to control glucosinolate chain length through a simple evolutionary pattern across the lineage (Abrahams et al., 2020). A broader phylogenetic analysis across 40 species revealed a more complex scenario where seven MAMs evolved through recurrent and convergent events involving tandem duplications, neofunctionalization, and independent gene losses (Abrahams et al., 2020). This highlights the need for comprehensive phylogenetic and functional studies to fully understand the evolution of specialized metabolism within plant families.

To develop a deeper understanding of how specialized metabolism evolves within a family, we focused on the variation controlling production of 2-hydroxy-but-3-enyl (2HB3) glucosinolate. Metabolic analysis across the *Brassicaceae* suggests that there are potentially extensive changes in GSL-OH including both presence/absence and enantiomeric variation in the reaction. *A. thaliana* accumulates a fixed ratio of 2R and 2S enantiomers of 2HB3, *Brassica napus* makes solely 2R, and closely related *Crambe abyssinica* make the 2S enantiomer. The side-chain modification to make this compound depends on two enzymes, GSL-OH and AOP2. The AOP2 enzyme converts methylsulfinyl glucosinolates to alkenyl glucosinolates like 4-methylsulfinylbutyl to but-3-enyl glucosinolate, which the epistatic GSL-OH then converts to 2-hydroxy-but-3-enyl (2HB3) glucosinolate (Figure 1). As such, studying the variation in 2HB3 glucosinolate across the *Brassicaceae* and how this connects to GSL-OH and AOP2 variation allows us to test the interplay of epistasis between loci, gene duplication, gene loss and enzymatic variation in controlling glucosinolate diversity.

**Fig. 1:**
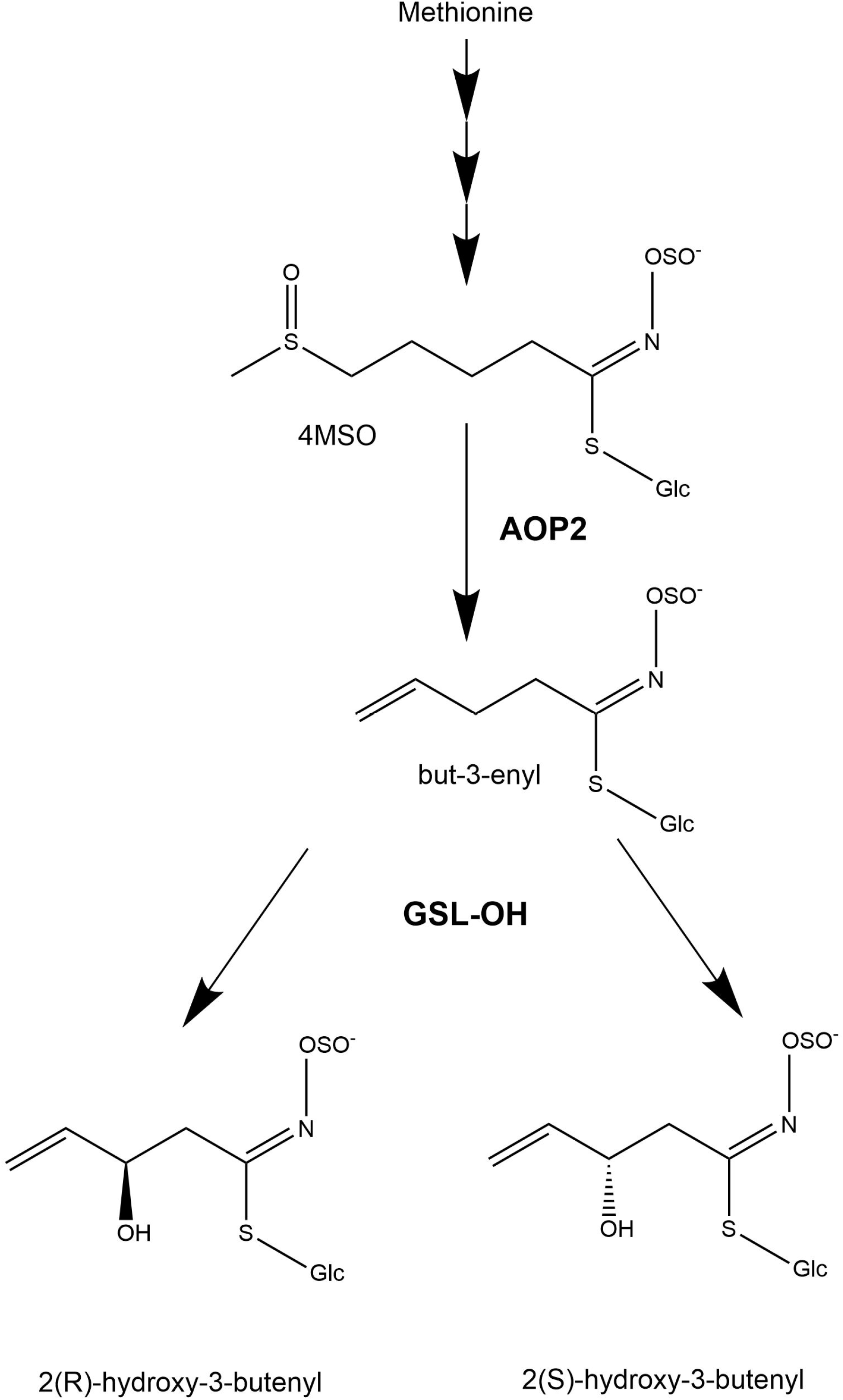
2HB3 biosynthetic pathway. In *A. thaliana* AOP2 converts but-3-enyl to 4-MSO. GSL-O catalyzes the conversion of but-3-enyl to both enantiomeric forms of 2-hydroxy-but-3-enyl. A functional AOP2 and GSL-OH expression are required for the synthesis of 2HB3.

To empirically map how specialized metabolite diversity is evolving across shorter evolutionary time frames, we tested how the AOP2 and GSL-OH enzymatic steps evolved across the Brassicaceae. We leveraged genomic and chemical data from 46 Brassicaceae species and identified candidate GSL-OH and AOPs orthologs. Functional validation of 36 putative orthologs was conducted via complementation assays expressing these putative orthologs in stable transgenic lines of an *A. thaliana* accession that only produces the but-3-enyl precursor due to a knockout mutation in GSL-OH. This enabled us to map the evolution of GSL-OH function across the *Brassicaceae* and investigate potential co-evolutionary connections with the AOP2 enzyme. From here, we combined the gene and metabolite data with the phylogenetic data to discern patterns in evolution across the *Brassicaceae* and estimate times of origin for these genes, identifying novel roles for segmental and distal duplication along with potential convergent shifts in stereofunctionality.

## RESULTS

### 2HB3 variation in the *Brassicaceae*

To map chemical diversity, we collated the available literature on GSL-OH presence and functionality across the *Brassicaceae* family to provide a phylogenetic context of this diversity. We identified 49 *Brassicaceae* species with published genomes and refined species phylogeny (Hendriks et al., 2023). We then collected glucosinolate content information from published datasets (Fahey et al., 2001; Brown, J. and Morra, M.J., 2005) (Supplemental Table 1). We used the presence of 2HB3 to infer the existence of a GSL-OH ortholog. Similarly, if 2HB3 or but-3-enyl were reported to be present in the species, we assumed the presence of a functional AOP2. If no alkenyl glucosinolate was identified in the species, this suggests an absence of AOP2 meaning it is not possible to infer GSL-OH and it was listed as not identifiable. This allowed us to estimate GSL-OH and AOP2/3 activity across the Brassicaceae family (Figure 2). We were able to obtain glucosinolate information on 42 of the species, showing that 17 species contain 2HB3, although enantiomer specificity is not reported for the majority. Of the 25 species not containing 2HB3, 14 lacked but-3-enyl and were inferred to lack AOP enzymatic activity. Eight species contained but-3-enyl, but no 2HB3, indicating a loss of GSL-OH activity. Mapping these events on a broader species tree shows extensive independent presence-absence variation across the species for all major lineages (Figure 2). However, there did appear to be more absence of AOP2 and thus the GSL-OH product within the Camelinodae tribe (including *A. thaliana*) while the Brassicodae supertribe had more widespread maintenance of GSL-OH and AOP2 activities (Figure 2). For GSL-OH, the metabolic presence-absence variation suggests at least three independent losses within the 40 MYA divergence time amongst these species. To empirically validate these gene presence/absence predictions we proceeded to use the genome sequences to identify AOP2/3 and GSL-OH genes and how it maps to GSL-OH activity and 2HB3 distribution across the *Brassicaceae*.

**Fig. 2:**
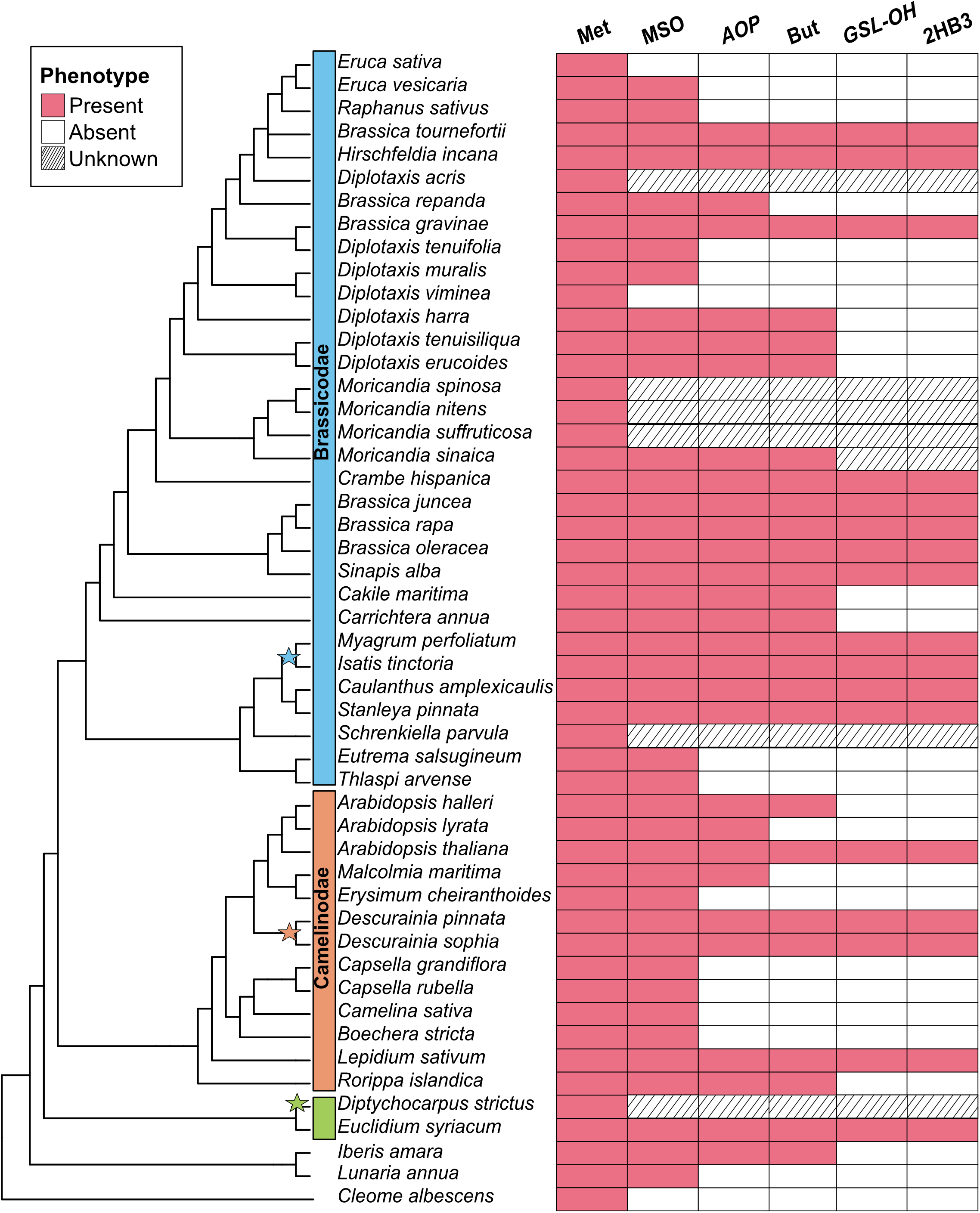
Phylogeny of 49 Brassicaceae species and C/eome *albescens* as outgroup, adapted from Hendriks et al. (2023). Left stripes indicate supertribes: Brassicodae (blue, lin II), Camelinodae (orange, lin I), and Hesperodae (green, lin Ill); unclassified taxa lack stripes. Right panel shows glucosinolate traits (multiple sources; Supplementary Table 1). Met represents methionine-derived glucosinolates, MSO stands for methylsulfinyl, AOP is for AOP2 that synthesizes but-3-enyl, and GSL-OH which converts the latter to 2-hydroxy-but-3-enyl. AOP and GSL-OH is inferred from metabolite presence. Stars denote potential independent whole-genome duplications per Mabry et al. (2020).

### AOP2/3 gene phylogeny

To systematically assess the role of AOP2 gene variation in controlling 2HB3 accumulation across the *Brassicaceae* we identified putative AOP2 orthologues we reduced the number of species to further analyze to 46 publicly available genomes of this family and *Cleome albescens*, a member of the sister family *Cleomaceae*, as an outgroup. Three species; *Brassica tournefortii, Arabidopsis halleri*, and *Brassica gravinae*, were discarded to reduce the number of potential candidate genes to test. *A. halleri* is known to lack GSL-OH (Windsor et al., 2005) and is similar to *A. lyrata*. Given the over representation of Brassica spp, *B. tournefortii* and *B. gravinae* were not further studied. Using the genomic sequences, we identified AOP2 homologues, performed sequence alignment, and constructed a phylogenetic tree using a previously developed pipeline (Steinbrenner, 2024). This tree showed the AOP2 and its tandem duplicate, AOP3, form a monophyletic clade with all alkenyl glucosinolate producing species having a AOP2 gene (Supplementary Figure. S1. and S2., Supplementary Table. S1.). The AOP3 tandem duplicate produces hydroxyalkyl glucosinolates that are only present in *Arabidopsis thaliana* and *Malcolmia maritima* and this is reflected by both species having a monophyletic AOP3 copy. Twelve of the species do not contain any sequence within the AOP2/3 clade: *Raphanus sativus, Lepidium sativum, Eutrema salsugineum, Erysimum cheiranthoides, Eruca vesicaria, Eruca sativa, Diplotaxis viminea, Diplotaxis tenuifolia, Diplotaxis muralis, Diplotaxis acris, Caulanthus amplexicaulis*, and *Brassica oleracea var. capitata*. None of these species had alkenyl nor 2HB3 glucosinolate. This suggests that presence/absence variation across the phylogenetic tree for alkenyl-derived glucosinolate production is mediated by loss of the AOP2 gene.

### GSL-OH gene phylogeny

To begin a systematic assessment of GSL-OH gene distribution across the *Brassicaceae* we used the same 46 publicly available genomes. While the *Brassicaceae* family contains five major supertribes, the Brassicodae and Camelinodae are overrepresented in the genomes currently available (Figure 2). As before, we used the previously developed blast-align-tree pipeline (Steinbrenner, 2024) to construct a phylogenetic tree to identify all potential GSL-OH orthologues and paralogues. The resulting phylogeny contained a total of 247 genes from 46 species (Supplementary Figure. S3). Parsing the tree by the three *Brassicaceae* supertribes suggested at least four larger gene families within the tree.

We found 51 genes within a monophyletic clade that contained the *A. thaliana* GSL-OH (AT2G25450). Two closely related monophyletic clades were represented by the *A. thaliana* genes AT2G30830 and AT2G30840. Neither AT2G30830 nor AT2G30840 are required for *A. thaliana* GSL-OH activity. To validate that these were not GSL-OH encoding enzymes, we performed a complementation assay in an *A. thaliana* accession that accumulates solely but-3-enyl precursors due to a natural GSL-OH knockout. The functional GSL-OH/AT2G25450 complemented the ability to synthesize 2HB3 glucosinolate in 70 total independent transgenics while, neither AT2G30830 nor AT2G30840 complemented 2-hydroxy-but-3-enyl production (minimum 15 independent transgenic events) supporting that these closely related clades are not functional GSL-OH enzymes. Both AT2G30830 and AT2G30840 have no stop codons, contain all necessary active site residues for a 2-oxoacid dependent dioxygenase, and are expressed in the transgenics suggesting they likely conduct a different enzymatic reaction. Thus, we hypothesized that the monophyletic clade of *Brassicaceae* genes containing the *A. thaliana* GSL-OH, AT2G25450, is likely the functional GSL-OH containing clade (Figure 3). We proceeded to functionally validate 36 of the 51 genes in this clade, representing all three sampled supertribes, using the above *A. thaliana* stable transgenic assay to test this hypothesis (Figure 3 and Supplemental Table. S3.); reasoning for the 36 selected can be found in the text and Supplemental Table 3. These results are discussed below by phylogenetic grouping.

**Fig. 3:**
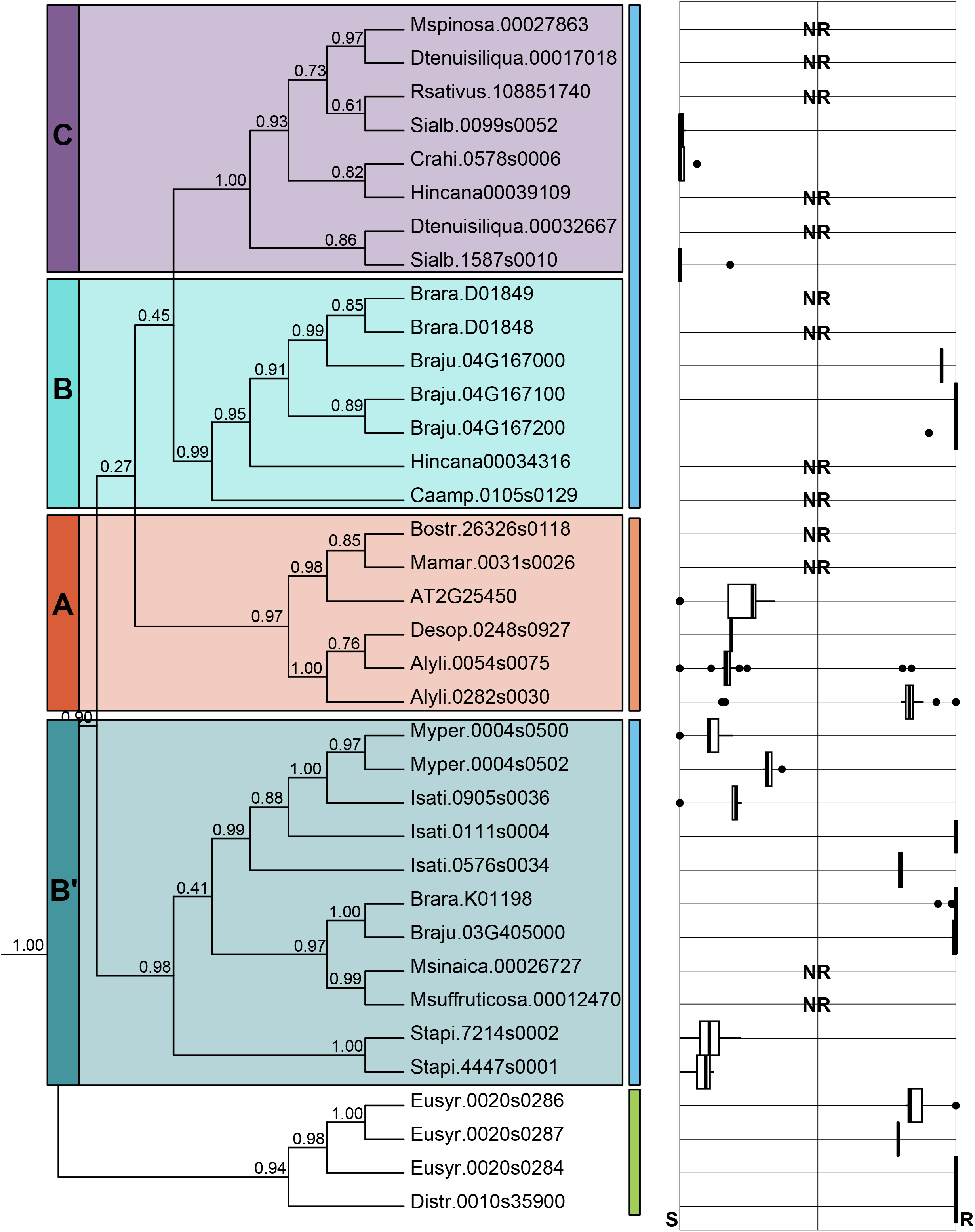
GSL-OH phylogenetic tree constructed with 46 Brassicaceae genomes using the Steinbrenner lab’s blast-align-tree pipeline. Selected GSL-OH orthologs were functionally validated. Box plots indicate enantiomer preference calculated as R/R+S. Nonfunctional genes lack a boxplot, instead having NR for no reaction.

#### Camelinodae supertribe functional validation

Given the phylogenetic proximity to the validated *A. thaliana* GSL-OH (GSL-OH A; Figure 3), we began functional validation tests by focusing on all orthologs from species in the Camelinodae supertribe. Members from the super tribe Camelinodae with GSL-OH orthologs grouped in a single clade. We included genomes from twelve Camelinodae species; seven species had no reported 2HB3 accumulation and accordingly lacked candidate genes in the GSL-OH clade. Their genome sequences also had no identifiable GSL-OH fragments, suggesting the gene has been fully deleted from all eight species. To further assess GSL-OH gene loss in Camelinodae, we used the high-quality genomes from the *A. thaliana* relatives, *Arabidopsis lyrata* and *halleri*. Both species have been documented to lack 2HB3 accumulation and *A. halleri* also lacks AOP2 (Stolpe et al., 2017). Neither species had any GSL-OH homolog and an investigation of the syntenic region showed no residual fragments of the gene indicating a complete loss. As a control, we selected the closest homologous gene in *A. lyrata*, AL4G25790, that resides in the AT2G30840 non-GSL-OH family of the tree and tested its potential GSL-OH activity. The gene showed no ability to rescue accumulation of 2HB3 in *A. thaliana* transgenics (10 T1 events). This gene had no premature stop codons, all necessary active site residues and was expressed in the transgenics suggesting it is not a GSL-OH-like enzyme and likely performs another function. This suggests that the absence of 2HB3 accumulation in seven of the twelve Camelinodae species is caused by whole gene deletions of GSL-OH.

To confirm the function of the six identified GSL-OH Camelinodae supertribe genes, we tested AT2G25450, Alyli.0054s0075, Alyli.0282s0030, Desop.0248s0927, Mamar.0031s0026, and Bostr.26326s0118. The genome sequenced as *Alyssum linifolium* (Alyli) has now been identified as *Descurainia pinnata* (Abrahams, R.S., et al., 2020). These genes were cloned and transformed into the *A. thaliana* accession Sha-1, which does not naturally produce 2HB3 as described above. As expected, AT2G25450 complemented the production of 2HB3 in all ten independent T1 plants and T2 progeny. Alyli.0054s0075, Alyli.0282s0030 and Desop.0248s0927 led to the accumulation of 2HB3 in all T1 and T2 transgenic leaves (minimum of 10 independent T1 per gene). Interestingly, the two *Descurainia pinnata* genes, accumulated different enantiomeric ratios of 2HB3, Alyli.0054s0075 produces a ratio of 1R:5S 2HB3 while Alyli.0282s0030 produces the opposite ratio of 5R:1S (Figure 3). These two genes differ in only 15 amino acids, suggesting that at least some of these changes are responsible for enantiomeric variation in GSL-OH.

*Boechera stricta* and *Malcolmia maritima* both have a GSL-OH orthologue but do not accumulate 2HB3 glucosinolate. In agreement with this, neither gene could rescue the accumulation of 2HB3 in any tested *A. thaliana* T1 or T2 generation (minimum of 10 independent T1s) (Figure 3). Neither species has a functional AOP2, nor does it make the precursor alkenyl glucosinolate. This suggests that the losses of AOP2 and the epistatic GSL-OH occurred in a relatively close time frame. Supporting this is a survey of the *M. maritima* GSL-OH protein sequence showing a premature stop codon similar to the one seen in some *A. thaliana* accessions with a non-functional GSL-OH (Supplementary Figure 5). In contrast, the *B. stricta* GSL-OH gene, Bostr.26326s0118.1, had no obvious genetic lesions, contained all active site amino acids required to be a functional 2-oxoacid-dependent dioxygenase (2-ODD)(Supplementary Table 3), and the RNA accumulated in all transgenic lines, yet did not enable 2HB3 production. Thus, it is possible that this gene has evolved to carry out a different reaction, allowing the gene sequence to be maintained in the absence of the but-3-enyl precursor or GSL-OH activity. This shows that GSL-OH function is lost via both whole gene deletions and nonsense mutations.

#### Brassicodae supertribe functional validation

We next proceeded to investigate the putative GSL-OH orthologues in the supertribe Brassicodae. We used 30 genomes from this supertribe, most of which belonged to the Brassiceae tribe. Eleven of these species had no 2HB3 glucosinolate nor a GSL-OH gene, indicating complete gene losses (Figures 2 and 3). Seventeen of the species had a gene that was positioned within the GSL-OH clade (Figures 2 and 3). The distribution of the Brassicodae GSL-OH orthologs across the gene tree indicated that there are three major clades. These clades form a polytomy with the clade containing Camelinodae species, suggesting they rapidly arose near the divergence of the Camelinodae and Brassicodae supertribes (Figure 3). We refer to these clades as Subclade C, Subclade B, and Subclade B’. Subclade *C* contains genes solely from species in the Brassiceae tribe. The next subclade, *B* subclade, contains sequences from Ca*ulanthus amplexicaulis*, and a few members of the Brassiceae tribe. The third subclade, *B’* subclade, appears to have originated prior to the split between the Brassicodae and Camelinodae. Subclade *B’* predominantly contains Brassicodae species that are not in the Brassiceae tribe along with two Brassiceae. Most species contain a GSL-OH in only one subclade while no species have genes in all three subclades. Three species, *Hirschfeldia incana, B. juncea*, and *B. rapa*, have genes in two of the subclades.

Although Brassiceae tribe is marked by a shared whole-genome triplication (WGT) event, and we identified three GSL-OH-like clades that contain Brassiceae species, it does not appear that the Brassiceae WGT event contributed to the origin of any of the three clades, nor did it expand them. Subclades B and B’ both contain species that diverged prior to the WGT event, indicating their origins predate the event. Additionally, most genes appear to have rapidly lost their WGT-derived duplicates following the triplication, as all homologs from tribe *Brassiceae* species in these clades are either single-copy or tandem duplicates (Figure 3). Subclade C only contains species in the tribe Brassiceae, but only one species, *Sinapis alba*, contains more than one copy suggesting species in this clade also reduced to single copy after the WGT event (Figure 3 and Supplementary Figure S4). To resolve potential functional relationships within the gene tree, of the 41 Brassicodae genes in the GSL-OH family we functionally validated 36 (Supplementary Table 1).

##### I. *B* subclade

The *B* subclade was found in the fewest species with sequences in only 5 of 46 total (Supplementary Figure 4 and Supplementary Table 3) Brassicaceae genomes. This included two tandem genes in *B. rapa* previously reported as potential GSL-OH genes; Brara.D01848 and Brara.D01849. Neither gene was functional in our validation assays possibly due to both sequences missing 50 amino acids at the C terminus, two of which are involved in 2ODD active sites. Checking the annotated genome confirmed the early stop codons in these genes. *B. juncea* is a natural allotetraploid of *B. rapa* and *B. nigra* and contains three genes that group with the non-functional *B. rapa* genes: Braju.04G167000, Braju.04G167100, and Braju.04G167200. These proteins are full-length and were all functional (Figure 3 and Supplemental Table 3). Of the remaining tested genes in the B subclade, only the *Caulanthus amplexicaulis* ortholog was weakly functional, with very low amounts of but-3-enyl being converted to 2HB3. The Hincana00034316 gene was non-functional and contained active site mutations (Supplementary Table 3).

##### II.*C* subclade

Subclade *C* contained GSL-OH candidates from a wide range of species, including ones not known to produce 2HB3. These sequences were found in only 8 of 46 total Brassicaceae species (Supplementary Figure 4 and Supplementary Table 3). Given the presence-absence-variation for the compound observed across species we decided to test them all and explore possible explanations for the PAV observed. Of the thirteen genes observed, eight were tested, and only three were functional; Crahi.0578s0006, Sialb.1587s0010, and Sialb.0099s0052. Querying the other ten genes showed that four genes had altered active site amino acids likely abolishing their function (Supplemental Table). This subclade also had independent changes in enzyme stereospecificity as all the functional genes in this subclade made predominantly S2HB3.

#### *III.B’* subclade

The B’ GSL-OH subclade had the fewest sequences within the Brassicae tribe, only *B. rapa* and *junceae*, and sequences in only X of Y total Brassicaceae genomes. This included the only functional *B. rapa* GSL-OH gene, Brara.K0119, that, along with the close homolog Braju.03G405000, synthesized the R enantiomer. In addition to these two Brassica sequences, the B’ subclade is the only subclade to have sequences from *Stanleya pinnata* and species in the tribe Isatideae of Brassicaceae: *Isatis tinctoria* and *Myagrum perfoliatum*. The *S. pinnata* and *M. perfoliatum* genes were all functional with a preference to make the S enantiomer (Figure and Supplemental Table). Interestingly, *I. tinctoria* represented the only species that contained a large expansion in the number of GSL-OH with *I. tinctoria* having nine GSL-OH homologous. This expansion in *I. tinctoria* and to a lesser extent *M. perfoliatum* maybe linked to a neopolyploidy event (Supplemental Figure 4 and 5) (Mabry et al., 2020). To test the function across this gene expansion we focused on Isati.0905s0036, Isati.0576s0034 and Isati.0111s0004.1. All three genes could rescue 2HB3 production, each with a distinct enantiomeric specificity (Figure 3).

### Genomic origin of key GSL-OH loci

The phylogenetic analysis of the GSL-OH homologs suggested that the gene family expanded before the split between the Camelinodae and Brassicodae to create four distinct loci reflected as subclades in the tree (Figure 3). To better understand how this occurred genomically, we used the syntenic relationship between the genes across the species. Comparative genomic analysis suggested that the locus likely first arose from a tandem array of 2ODDs that currently maps to chromosome 4 in *B. juncea* and represents the GSL-OH B locus (Figure 4). In the B locus, the functional GSL-OH is located between the neighboring non-GSL-OH 2ODDs (*AT2G30830* and *AT2G30840* in *Arabidopsis thaliana* and Braju.4G163000 and Braju.4G168000 in *B. juncea*). These flanking 2ODD genes are the most closely related genes to GSL-OH, suggesting that a local tandem duplication and neofunctionalization may have created the original GSL-OH (Figure 4). Local macrosynteny between the GSL-OH B and B’ loci indicates that the B’ locus within the Brassicodae arose by a segmental duplication of the B locus. In contrast, micro- and macrosynteny analysis of the GSL-OH C homologs in *S. alba*, and *H. incana* found no synteny to either the GSL-OH B or B’ loci. This suggests that GSL-OH C arose by a single gene distal duplication prior to the whole genome triplication in the Brassicaceae tribe.

**Fig. 4:**
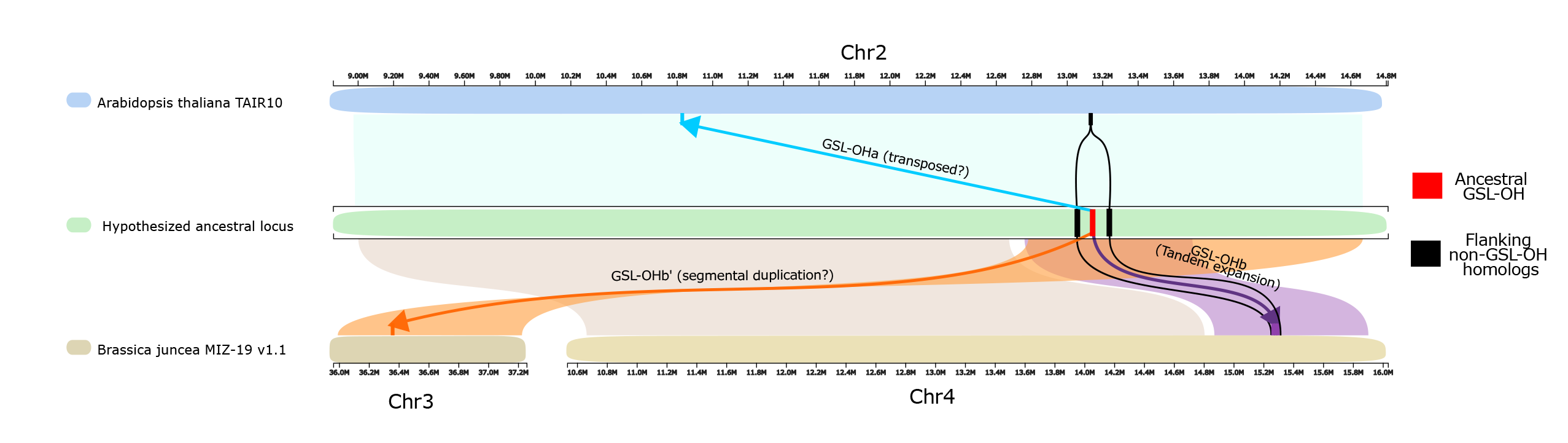
A model for the evolution of the GSL-OH locus across the Brassicaceae family. The physical positions of the Arabidopsis thaliana GSL-OH gene, called GSL-OHa (light blue), and the two closest homologs of GSL-OH, also of the 2-ODD enzyme family (black), are represented on the top chromosome in light blue. Colored ribbons represent macrosynteny to regions on two chromosomes of Brassica juncea show the position of two functional GSL-OH homologs, the tandemly duplicated GL-OHb genes (purple), GSL-OHb’ (orange), and homologs of the two closely related non-GSL-OH 2-ODD genes (black). here was no detectable macrosynteny between any of these regions and GSL-OHc homologs and so they are excluded from this figure. Based on the evolutionary relationships of these genes and the patterns of macrosynteny we present a hypothetical ancestral locus in green, where an ancestral GSL-OH was nested within the 2-ODD homologs, GSL-OHb retains the ancestral position, GSL-OHb’ arose by a segmental duplication or similar mechanism, and GSL-OHa was transposed to the current location 2MB upstream of the non-GSL-OH homologs or some similar mechanism. B) The evolutionary relationships of these GSL-OHhomologs is represented in the simplified phylogeny on the right.

Investigating synteny within the Camelinodae GSL-OH A locus shows that the GSL-OH B and B’ regions from the Brassicodae maps to an *A. thaliana* region on chromosome 2 from ∼12.8Mbp to 13.7Mbp. Within these are the two *A. thaliana* closest GSL-OH 2ODD relatives, AT2G30830 and AT2G30840 that flank GSL-OH in the B locus. However, unlike the GSL-OH B locus, this region does not contain the Arabidopsis GSL-OH. Instead, the GSL-OH gene is located approximately 2Mbp away in a genomic block showing no macro or microsynteny to any of the B, B’ or C genomic regions. This suggests that, like GSL-OH C, the GSL-OH A locus in the Camelinoidae arose via an independent distal gene movement out of the neighboring GSL-OH B genomic position. This shows that the evolution of the gene family within these species very likely involved segmental duplications, distal duplications, and distal gene movement from an ancestral locus that arose prior to Camelinodae and Brassicodae split.

### Querying for potential differences in enantiomers biological role

Across all four subclades, GSL-OH enzymes showed diversity in the specific enantiomeric mix that independently and convergently evolved across the phylogenetic tree. This raises the question of whether this variation in stereochemistry is functionally relevant for plant-biotic interactions or is largely neutral. Previous studies (Hansen, et al., 2008) have shown the presence of the *A. thaliana* GSL-OH, which makes a 1R:3S 2HB3 mix, and reduces Trichopulsia *ni* herbivory. To test if there is a potential difference in biological functions between the 2HB3 enantiomers, we used multiple independent and stable transgenic lines expressing either *Descurainia pinnata* Alyli.0054s0075 (Alyli54) or Alyli.0282s0030 (Alyli28) that make either the S or R enantiomer, respectively (Figure 3). We tested biotic interactions in these lines in combination with the parental CVI accession that produces the precursor glucosinolate, but-3-enyl. For each transgene, three independent transgenic lines with similar 2HB3 and total glucosinolate levels were chosen. These lines were tested for both *T. ni* herbivory and *Botrytis cinerea* fungal resistance. In *B cinerea* detached leaf assays, Alyli54 (making mainly S) had significantly larger fungal lesions on average across six different botrytis strains than did either the transgenic lines expressing Allyl28 (making mainly R) or the control lines accumulating the precursor but-3-enyl (Figure 5A). This suggests that R2HB3 is less effective at providing resistance to multiple *B. cinerea* isolates than but-3-enyl and S2HB3. In contrast, *T. ni* choice tests showed a significant preference for consuming lines expressing Alyli28 (R2HB3) over Alyli54 (S2HB3), with 90% of choice trials having more R than S eaten (Figure 5B). Interestingly, the choices were altered when compared to the but-3-enyl control, with R being equivalent to the control and the control being preferred over the S. This suggests that the enantiomers and precursor have differential effects depending on the biotic attacker being tested and that the differences could be non-neutral. Broader more systemic assays will be needed to assess the breadth and types of interactions that are altered by the enantiomeric differences.

**Fig 5:**
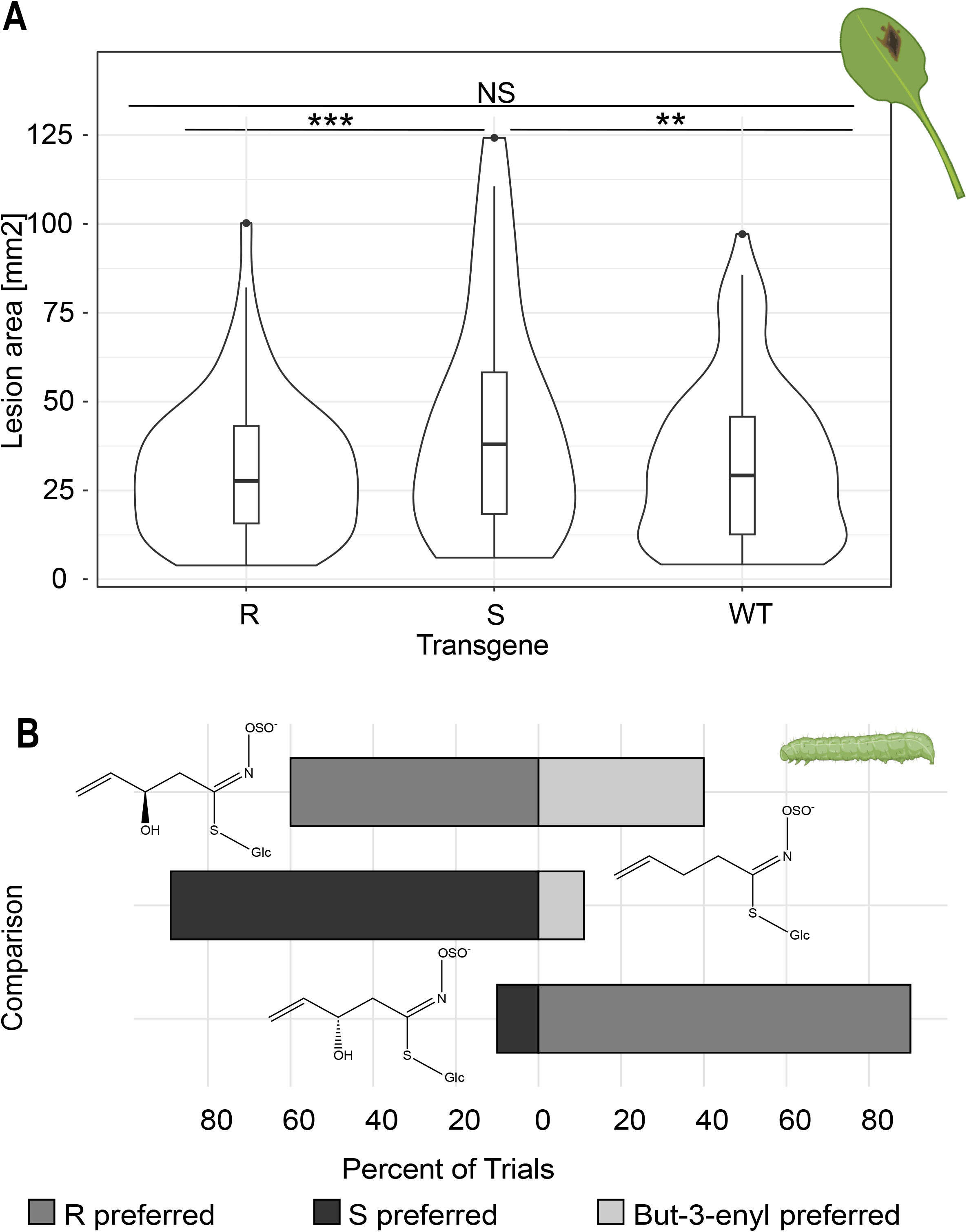
Effect of enantiomer variability in B. cinerea lesion size and T. ni herbivory.

## DISCUSSION

Using the genome sequences of 46 Brassicaceae species in conjunction with empirical validation allowed us to chart the evolution of the GSL-OH enzyme throughout the Brassicaceae family. This showed independent and convergent presence-absence and enantiomeric variation across the major *Brassicaceae* lineages studied. Genome sequences showed that a majority of this presence-absence variation is the result of recurrent complete gene loss of GSL-OH (Figures 2 and 3). We also identified a number of existing pseudogenes that have either active site missense or nonsense mutations (Figure 3 and Supplemental Table 3). Genomic synteny analysis showed that in the Brassicodae, this is occurring across a background with three distinct loci created by segmental and distal gene duplications that unequally partitioned across the tribe. In the Brassicodae species that had a whole genome triplication, this would have created a potential of nine gene copies, yet none of the Brassicoidae had more than 3 copies while seventeen had lost all 9 entirely. Species that contain a functional GSL-OH show evidence of independent changes in function whereby GSL-OH genes shift between making mainly the R or S enantiomers. These enantiomers had differential activity in two different biotic resistance assays to a fungus and insect. This shows that simple presence/absence phenotypic variation within a metabolic trait is caused by a highly complex set of genomic processes enabling this variation.

### Duplication events in gene evolution

Evolution of specialized metabolite genes or gene families is typically associated with tandem gene or whole genome duplication. In contrast, GSL-OH showed the importance of gene loss in conjunction with distal and segmental gene duplication in shaping gene family evolution (Figure 4). The *Brassicaceae* genomes did reveal tandem gene expansion/contraction, such as the expansion within *Isatis* of GSL-OH B’ to 9 copies. However, synteny analysis suggested that the primary driver of GSL-OH expansion/evolution was distal and segmental duplications that were not connected to whole genome duplication events. We hypothesize that the original GSL-OH arose within the *B* position as a tandem duplicate of one of the neighboring 2-ODDs. This then underwent a segmental duplication to create the *B’*. In the Camelinodae, the *B’* gene was lost and the B locus distally moved to a non-syntenic position, creating the A locus that is only present in the Camelinodae. In the Brassicodae, a further distal duplication created the C locus such that B, B’ and C were all present prior to the ancient whole genome triplication event in the Brassiceae tribe. Distal and segmental duplications have also been found creating variation in other specialized metabolite genes such as the origin of a novel MAM within the Brassicas and a novel CYP79F within *Boechera* (Abrahams, R.S., et al., 2020; Prasad et al., 2012; Schranz et al., 2009). Broader surveys for distal and segmental duplications in phylogenetically informed species are needed to ascertain if this is a broader feature of plant specialized metabolite evolution.

### GSL-OH Protein function variation

In addition to gene loss, our experimental phylo-functional approach identified an unexpected ability to shift enantiomeric specificity such that some GSL-OH proteins made predominantly the R, the S, or a mix of R/S enantiomers (Figure 3). The enantiomeric specificity rapidly changed both between and within species. At its extreme, *I. tinctoria* had tandem gene duplications of a single locus creating nine copies of GSL-OH that could be grouped into three structurally similar proteins. Each group synthesized one of three combinations: predominantly R, predominantly S, or only R. This enantiomeric shift occurred across all four GSL-OH loci, such as a pair of tandem GSL-OH homologs in *D. pinnata* where one copy synthesized a 1R:5S ratio and the other 5R:1S. A correlation of active site polymorphisms with enantiomeric specificity found no connections suggesting that the shift may be created by altering the general shape of the active site pocket. This could shift the angle of oxygen attack on the butenyl glucosinolate precursor. Further work is needed to ascertain how enantiomeric specificity is generated.

The enantiomeric variation raises the question of whether stereospecificity has any biological relevance. Glucosinolates are often most active when catabolized to the isothiocyanate with the 2HB3 isothiocyanate spontaneously cyclizing depending on the enantiomer to either epi-goitrin(R) or goitrin(S). In human studies, goitrin affects iodine uptake by the thyroid while epi-goitrin has no effect. Previous data has shown that the Arabidopsis GSL-OH leads to more resistance to the herbivore *Trichoplusia ni but* could not test if the effect depended on the enantiomer (Hansen et al., 2008). Using the *D. pinnata* genes, we could show that the S enantiomer of 2HB3 alters resistance to a collection of *B. cinerea* isolates while the R enantiomer has no effect. In contrast, *T. ni* choice tests showed a significant preference for consuming lines primarily accumulating R over S, with 90% of choice trials having more R than S eaten (Figure 5B). Interestingly, the choices were altered when comparing to the but-3-enyl control with R being equivalent to the control and the control being preferred over the S. This suggests that the enantiomers have differential effects depending on the biotic attacker being tested. Fully understanding the potential biological consequences of this variation would require systemic assays taking these genotypes and assaying them against biotic and abiotic stresses both in the field and the laboratory.

## Materials and methods

### Visualization of species phylogeny and glucosinolate distribution

To assess our current understanding of GSL-OH across the Brassicaceae we kindly received a recently published species phylogenetic tree (Hendriks et al., 2023). We narrowed it down to species that have publicly available transcriptomes and produce 2HB3 or if a close relative synthesizes it. For example, in lieu of *Capsella rubella*, which we studied, the tree shows *Capsella bursa-pastori*. The glucosinolate profiles were primarily obtained from (Fahey et al., 2001) and additional literature as needed. We then incorporated the glucosinolate information to ascertain if species likely contain AOP2 activity, but-3-enyl GLS, GSL-OH activity, and or 2HB3 GLS.

### Building AOP2/3 and GSL-OH gene phylogeny

To construct the AOP2/3 and GSL-OH gene phylogenies we implemented the ‘blast-align-tree’ pipeline developed by the Steinbrenner laboratory at the University of Washington (Steinbrenner A. 2024. steinbrennerlab/blast-align-tree: BAT v0.1.1 (v0.1.1)) We obtained publicly available Brassicaceae coding sequences via Phytozome13 and those published in (Guerreiro et al., 2023). Using these local genomes, we implemented the pipeline with the default parameter on the coding sequences. However, to account for genome duplication events we altered the number of genes called from five to ten for certain species. The tree was redrawn and visually inspected to ensure that the new genes being identified were external to the gene tree of interest. If all new genes were internal to the clade of interest, we increased the number of genes to pull from a species until no new internal genes were identified. This ensured we obtained all possible orthologues. The resulting tree was visualized using the ‘ggtree’ package in R studio (Yu et al., 2017).

### Cloning of candidate GSL-OH genes

From the phylogenetic analysis, we identified the most closely related GSL-OH/AT2G25450 candidates within each species based on a shared evolutionary pattern. The outgroup clades were defined using the AT2G30840/30 genes unable to catalyze the GSL-OH reaction. This was done regardless of the reported presence of 2HB3 (Fahey et al., 2001). All candidate genes were synthesized by Azenta life sciences or Twist Biosciences. The expression vector selected was pFAST-R05 (Shimada et al., 2010) acquired from the VIB-UGENT center from plant systems biology. The genes generated by Azenta were cloned into the pFAST-R05 plasmid with Gateway cloning and pDONR221 as the donor vector. Twist biosciences genes were directly incorporated into the plasmid pFAST-R05. These were then transformed into *Agrobacterium tumefaciens*.

### Plants transformation

The floral dip method of *Agrobacterium tumefaciens*-mediated transformation of A. thaliana was used for plasmid insertion (Zhang et al., 2006). The A. *thaliana* accession Shakdara-1 or Cvi-0 were used (Doody et al., 2022) as they produce but3-enyl glucosinolate, the precursor metabolite needed for the GSL-OH reaction, but they do not synthesize the GSL-OH products, (R, S) 2-hydroxy-but-3-enyl. Most transformations were in the Shakdara-1 accessions due to a higher transformation efficiency

### Transgenic plants segregation test

GSL-OH is functionally dominant allowing its function to be measured in T_1_ seeds (Hansen et al., 2008). To measure the ability of the transformed enzymes to conduct the expected reactions, T_1_ Seeds emitting red fluorescence when placed under a 532nm laser attached to a camera lens were selected. These seeds were sown on soil and after rosette formation 40mg of tissue were collected for glucosinolate analysis extraction. The plants were allowed to self and T_2_ seeds were collected. These were also screened to confirm the potential enzyme activity. For each gene being tested, at least 5 independent T_1_ seeds were tested by HPLC and reconfirmed as T_2_.

### Glucosinolate extraction and analysis

To estimate GSL-OH function in all genotypes, GSLs were measured as previously described (Kliebenstein et al., 2001). Briefly, the leaf material was placed in 400 μL of 90% methanol. Samples were homogenized for 3 min in a paint shaker, centrifuged, and the supernatants were transferred to a 96-well filter plate with DEAE sephadex. The filter plate with DEAE sephadex was washed with water, 90% methanol and water again. The sephadex-bound GSLs were eluted after an overnight incubation with 110 μL of sulfatase. Individual desulfo-GSLs within each sample were separated and detected by HPLC-DAD, identified, quantified by comparison to standard curves from purified compounds. In this protocol, the R and S enantiomers of 2HB3 separate by about 2 minutes (Hansen et al., 2008).

### Transgene presence verification

To verify that the GS-OH gene was being expressed in the independent T1s, leaf tissue was collected from at least three independent T1s per transgene and extracted RNA using Monarch® Total RNA Miniprep Kit (T2010S). We then used OneTaq® One-Step RT-PCR Kit (E5315S) with gene specific primers designed to anneal at the exon-exon junction. RT-PCR product was run on a 2% agarose gel at 120 volts for 30 minutes and visualized to test if the mRNA was present. All transgenes expressed mRNA.

### Querying for active site mutations in non-functional GSL-OH genes

The protein structure and relevant amino acids of the different GSL-OH-like candidate genes were inspected to determine if GSL-OH homologues with no activity had key missense mutations responsible for the lack of function. The active site amino acids were previously defined from crystal structures (Wilmouth et al., 2002). They identified Tyr-217, Tyr142, Phe-144, Lys-213, His 232, Asp 234, His 288, Arg 298, Phe-304, and Glu-306 as the amino acids involved in the reaction within anthocyanidin synthase (AT4G22880) and this agreed with other 2ODD enzymes. All GSL-OH protein sequences were aligned using the ClustalOmega alignment in the msa R studio package with the default parameters (Bodenhofer et al., 2015). Files were exported and visualized in the desktop version of the software MEGA (Tamura et al., 2021). The corresponding active site amino acid or the lack of thereof in each functional and non-functional orthologue was determined from the alignment (Supplementary Table 3 and Supplementary Figure 5).

### Identifying syntenic relationships between genes

The evolutionary trajectory of the GSL-OH gene family was mapped using a multi-step approach necessitated by the level of presence/absence variation in this gene family across the species. To test the relationship of the Camelinodae GSL-OH vs the Brassicodae GSL-OH, by mapping macrosynteny between the *A. thaliana* TAIR 10 genome and *B. juncea ssp*. Integrifolia MIZ-19 genome v1.1 from a pre-compiled run of GENESPACE (Lovell et al. 2022) using the GENESPACE synteny viewer on Phytozome (Goodstein et al. 2012). This allowed focusing on the segments containing the identified *A. thaliana* GSL-OH and homologs Braju.3G405000, and Braju.4G167000. To better ascertain the relationship amongst *Brassicaceae* GSL-OH family members, the online CoGe platform was used (Lyons et al. 2008) to search for both micro- and macrosynteny between *S. alba* (Yang et al. 2023) compared to the GSL-OH gene from the *A. thaliana* TAIR 10 and *B. juncea* MIZ-19 genomes using the SynMap (Haug-Baltzell et al. 2017) and SynFind (Tang et al. 2015).

### *Botrytis cinerea* detached leaf assay and *Trichoplusia ni* choice test

Multiple independent transgenic lines of each of the Alyli.0054s0075 and Alyli.0282s0030 transgenes were used to test resistance to the necrotrophic fungi *B. cinerea*. These were selected given their stark difference in R vs S accumulation and equal levels of 2HB3 production. For this, homozygous transgenic T2 generation were grown for each gene alongside wildtype genotypes. For resistance assays, *B. cinerea* diverse strains were: 2.04.12, ROSE, B05.10, A517, KernB1, and KT. This allows assessing the consistency of the interaction across pathogen diversity. For each T1 transformation event, six T2 plants were selected, with six leaves chosen from each plant. Each leaf was inoculated with one of six fungal strains following the protocols from (Caseys et al., 2021). This resulted in 36 leaves per original transformation event, and this process was repeated for all four transformation events, as well as controls for each T2 plant. Pictures of the developing lesions were taken at 48-, 72- and 96-hours post-infection. The 72-hour time was selected for image analysis, and infection size digitally measured as described in (Caseys, et al., 2021). Statistical analysis was performed in R to obtain ANOVA values and corresponding figures.

For herbivory assays, six independent T2 transformants of Alyli.0054s0075 and Alyli.0282s0030 were used. Six leaves per plant were hole-punched for replicates, with each replicate consisting of two transgenic and two control leaves placed across one another on agar petri dishes. Three replicates per transformation event were used for a total of eighteen plates per transgene-wildtype grouping. Five *Trichoplusia ni* larvae were placed in the center of each replicate, and images were taken at 24 and 48 hours, with the 24-hour time point selected for analysis. Leaf consumption was quantified using GIMP, and the average percent consumed per gene and wildtype group within each plate was calculated for statistical analysis and figure generation in R.

