## Supplementary figures and images for "Convergence and constraint in glucosinolate evolution across the Brassicaceae"

### Supplementary figure 1

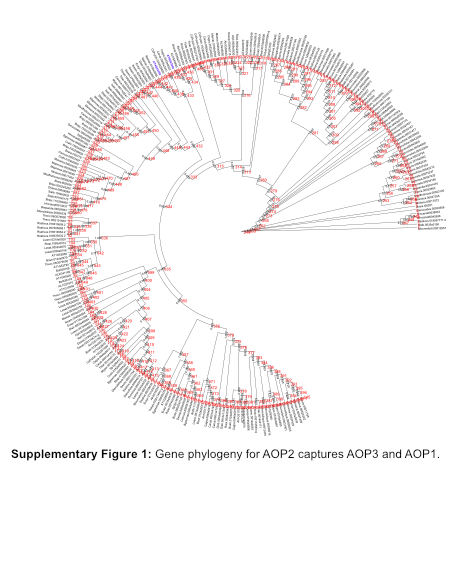

### Supplementary figure 2

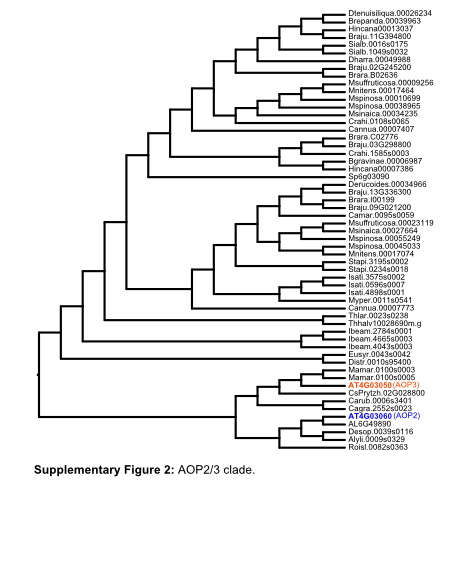

### Supplementary figure 3

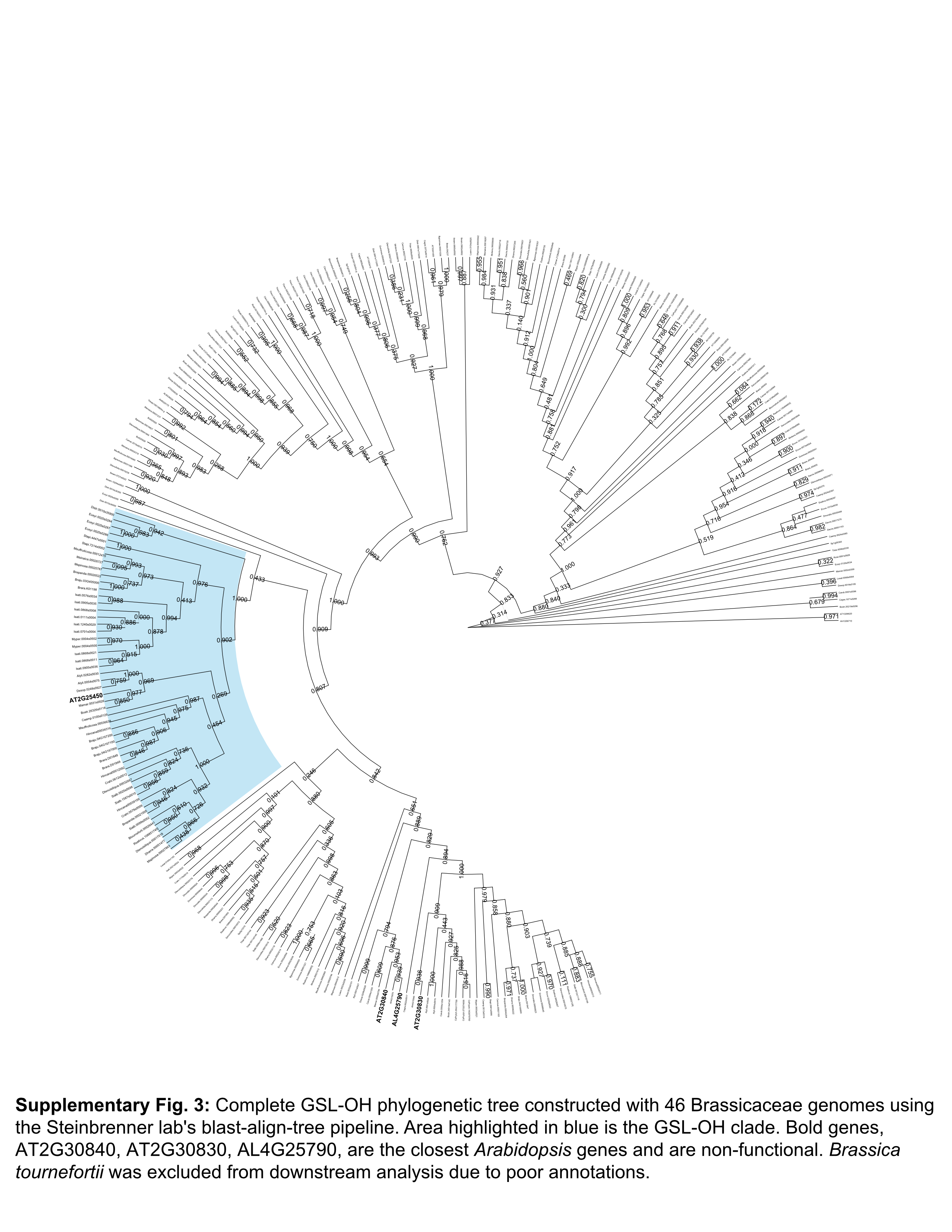

### Supplementary figure 4

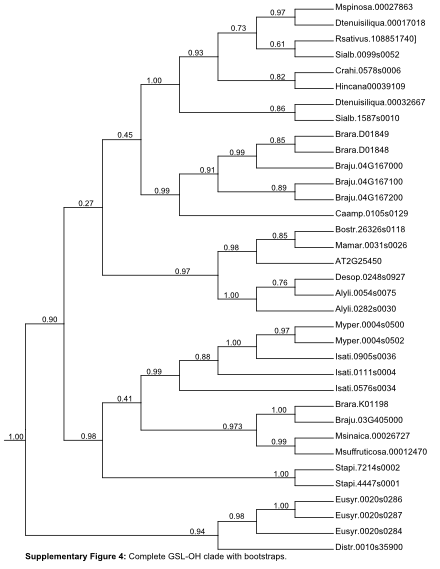

### Supplementary figure 5

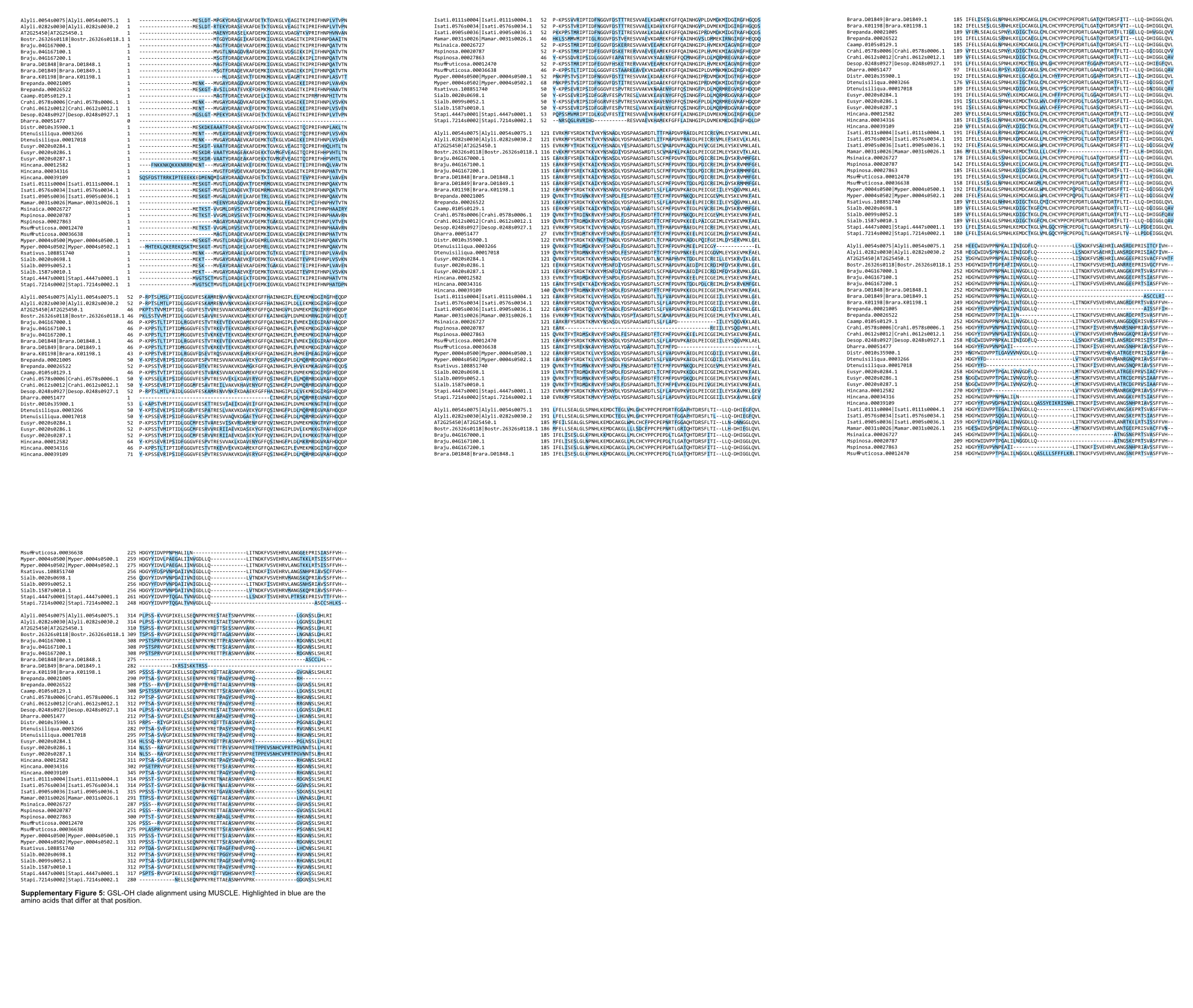

### Supplementary figure 6

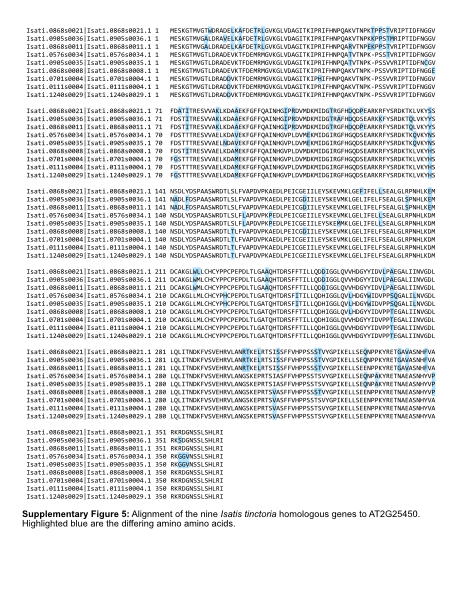
